# Early striatal hyperexcitability in an *in vitro* human striatal microcircuit model carrying the Parkinson’s *GBA-N370S* mutation

**DOI:** 10.1101/2023.03.01.530566

**Authors:** Quyen B. Do, Bryan Ng, Ricardo Marquez Gomez, Dayne Beccano-Kelly, Naroa Ibarra-Aizpura, Maria-Claudia Caiazza, Charmaine Lang, Jimena Baleriola, Nora Bengoa-Vergniory, Richard Wade-Martins

**Affiliations:** Oxford Parkinson’s Disease Centre and Department of Physiology, Anatomy and Genetics, University of Oxford, South Park Road, Oxford OX1 3QU, United Kingdom; Kavli Institute for Neuroscience Discovery, University of Oxford, Dorothy Crowfoot Hodgkin Building, South Park Road, Oxford OX1 3QU, United Kingdom; Aligning Science Across Parkinson’s (ASAP) Collaborative Research Network, Chevy Chase, MD, 20815, USA; Achucarro Basque Center for Neuroscience, Leioa, Spain; University of the Basque Country (UPV/EHU), Department of Neuroscience, Leioa, Spain; Ikerbasque - Basque Foundation for Science, Bilbao, Spain

## Abstract

Understanding medium spiny neuron (MSN) physiology is essential to understand motor impairments in Parkinson’s disease (PD) given the architecture of the basal ganglia. Here, we developed a custom three-chamber microfluidic platform and established a cortico-striato-nigral microcircuit recapitulating the striatal presynaptic triad *in vitro* using induced pluripotent stem cell (iPSC)-derived neurons. We found that, although cortical glutamatergic projections facilitated MSN synaptic activity, dopaminergic transmission was essential for excitability maturation of MSNs *in vitro*. Replacement of wild-type iPSC-dopamine neurons (iPSC-DaNs) in the striatal microcircuit with those carrying the PD-related *GBA-N370S* mutation induced early hyperexcitability in iPSC-MSNs through reduction of voltage-gated sodium and potassium intrinsic currents. Such deficits were resolved in aged cultures or with antagonism of protein kinase A activity in nigrostriatal iPSC-DaNs. Hence, our results highlight the unique utility of modelling striatal neurons in a modular and highly physiological circuit which is essential to reveal mechanistic insights of the loss of electrical functional integrity in the striata of *GBA1* PD patients.

## Introduction

The striatum is comprised of 90% medium-sized spiny neurons (MSNs) and represents the primary input nucleus of the basal ganglia ^1^. The striatum receives converging glutamatergic and dopaminergic inputs from the cortex and midbrain, respectively, forming a presynaptic triad critical for regulation of MSN functional activity. Striatal dysfunction plays a key role in the motor features of Parkinson’s disease (PD), although the exact underlying mechanisms remain under intense study with divergent findings ^2–4^. The consensus suggests that the characteristic preferential degeneration of substantia nigra *pars compacta* (SNc) dopaminergic neurons (DaNs) results in aberrant striatal activity patterns which subsequently alter the balanced output of the basal ganglia governing motor behaviour ^5^. However, the specific effects on the two parallel principle striatal pathways, namely the direct pathway and indirect pathway, are contentious. The classical premise suggested functionally opposed roles of direct and indirect pathway neurons in regulating action and their diametrical changes in firing rate in PD ^2^, although a more recent observation of co-activation of both pathways at the initiation of movement has challenged this view ^6^.

The advent of human induced pluripotent stem cell (iPSC) technology has revolutionized the ability to investigate the pathophysiology of neurological disease in human neurons, revealing a number of potential cellular mechanisms underpinning preferential degeneration of SNc DaNs ^7–12^. However, much less is known about how the presence of Parkinson’s mutations changes DaN function and how this in turn impacts MSN function. *In vitro* modelling of iPSC-derived human striatal neurons has developed from original striatal monocultures ^13–15^ to more recent three-dimensional cortico-striatal assembloids ^16^, however such models have not included dopaminergic neurons.

Microfluidic neuronal culture technology has tremendously assisted efforts in modelling and understanding biological systems, primarily enabling custom and precise control of neurite isolation ^17,18^ as well as orchestrated neuronal circuitry ^19,20^. In this work we have established a microfluidic platform comprising an all iPSC-derived striatal presynaptic microcircuit to recapitulate the endogenous connectivity of cortical and dopamine neurons onto MSNs. This setup is permissive to long-term cultures of up to 100 days and allows for the investigation of temporal changes in neuronal physiology. With this system we have dissected the importance of dual presynaptic glutamatergic and dopaminergic inputs in facilitating both electrophysiological maturation of *in vitro* MSNs as well as the early striatal pathology affected by the PD-related *GBA-N370S* genetic mutation.

## Results

### Generation of functional iPSC-derived medium spiny neurons (MSNs)

We first applied a defined cocktail of small molecules to recreate the temporal *in vivo* development of MSNs to reflect their telencephalic and lateral ganglionic eminence (LGE) lineage (Figure 1A). To induce forebrain development, we used modulators of SMAD, namely LDN and SB ^21^, and an inhibitor of the WNT signalling pathway (XAV) at the start of the differentiation ^22^. Activin A, which was independently reported to enhance dorsal striatal patterning ^14,16^, was introduced from DIV12-23 to promote LGE identity. Activin A from DIV 12 increased the expression of selected markers for LGE and MSN progenitors two days post treatment (Supplementary Figure 1A) as well as the proportion of post-mitotic neurons co-expressing DARRP32 and CTIP2 on DIV 35 (Supplementary Figure 1B). Temporal examination of transcript abundance of LGE and MSN progenitor markers using RT-qPCR suggested gradual patterning of iPSCs towards MSNs reminiscent of their *in vivo* developmental trajectory (Figure 1B-D). LGE-specific genes such as *DLX2, ASCL1*, and *GSX2* were enriched in early time-points and this enrichment was subsequently reduced in post-mitotic MSNs (Figure 1B). Meanwhile, expression of selected markers of MSN progenitors including *ISL1, DLX5, FOP2*, and *MEIS2* increased over time and remained abundant in post-mitotic MSN cultures (Figure 1C). *CTIP2, GAD61*, and *DARPP32*, which are indicative of post-mitotic MSNs, were virtually absent in early cultures and only subsequently expressed in great abundance from DIV 14 (Figure 1D). Tyrosine hydroxylase (*TH*) transcript was also detected in late cultures, implying the presence of potentially non-MSNs, TH-expressing neurons (Figure 1D). Notably, expression of *NKX2*.1, the medial ganglionic eminence marker, was barely detectable throughout the culture development (Figure 1B), indicating a strong LGE enrichment in the patterning.

**Figure 1:**
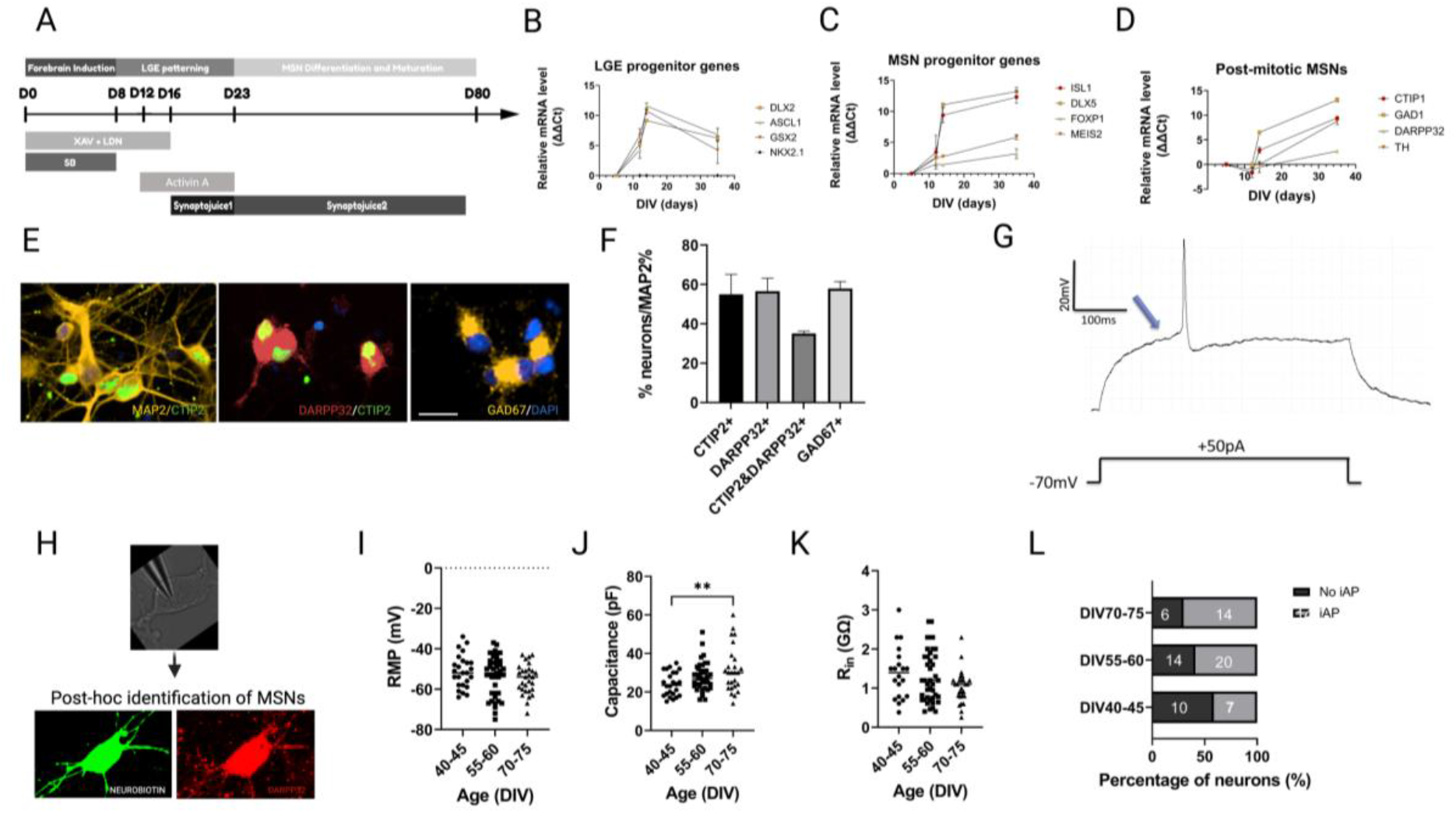
Generation and characterisation of iPSC-derived MSNs. (A) Differentiation schematic of medium spiny neurons (MSNs) from iPSC. (B-D) Temporal expression of selected markers of LGE progenitors (B), MSN progenitors (C) and post-mitotic MSNs (D) from DIV 0 till DIV 35. (E) Immunostaining for MAP2 (yellow), CTIP2 (green), DARPP32 (red), GAD67 (yellow) and DAPI (blue) and (F) quantification of their expression in percentage term at DIV 40-45. (G) Representative traces of evoked APs (iAP) showing slow-ramp depolarisation of MSNs. Blue arrow indicates delayed first spikes. (H) Whole-cell patch-clamp recording and subsequent post-hoc identification of neurobiotin-filled DARPP32+ neurons for electrophysiological analysis. (I-K) (I) Resting membrane potential (RMP), (J) cell capacitance, and (K) input resistance of iPSC-derived MSNs over time from DIV 40-75, n = 20-43 recording MSNs per condition, each dot represents a single neuron, One-way ANOVA corrected with Bonferroni post-hoc test, **p<0.01, ns p>0.05. (L) Number of MSNs that displayed evoked action potential upon current injection (iAP) and MSNs that did not (no iAP) over time. All data is presented as mean ± sem, N = 3-4 differentiation experiments of 3 iPS cell lines unless otherwise stated.

More than 50% of MAP2+ iPSC-MSNs at DIV 40 expressed CTIP2, DARPP32, or GAD67 and up to 37% co-expressed both canonical markers CTIP2 and DARPP32 (Figure 1E-F). We also observed expression of selected markers specific to either the direct- or indirect-pathway MSNs (Supplementary Figure 2A-B). Immunostaining confirmed expression of MSN-subtype specific neurotransmitter and neurotransmitter precursors including PENK, PDYN and SUBSTANCE P at DIV 40 (Supplementary Figure 2C). Quantification of GABAergic neurons co-expressing either PKEN, PDYN or SUBSTANCE P suggested the dominant presence of direct pathway MSNs intermingled with a modest population of indirect pathway MSNs in our iPSC-MSN cultures (Supplementary Figure 2D).

We next performed whole cell patch-clamping to examine functional maturation of DARPP32-positive iPSC-derived MSNs (Figure 1H). Our iPSC-derived MSNs displayed hyperpolarised resting membrane potential (DIV 40-45: −51.2 ± 1.62 mV, n = 25 neurons; DIV 55-60: −53.3 ± 1.47, n = 43 neurons; DIV 70-75: −55.1 ± 1.36 mV, n = 29 neurons), large cell capacitance (DIV 40-45: 24.0 ± 1.26 pA, n = 24 neurons; DIV 55-60: 28 ± 1.11 pA, n = 42 neurons; DIV 70-75: 31.6 ± 2.31 pA, n = 26 neurons), and large input resistance (R_in_) (DIV 40-45: 1.39 ± 0.15 GΩ, n = 20 neurons; DIV 55-60: 1.29 ± 0.10 GΩ, n = 41 neurons; DIV 70-75: 1.0 ± 0.08 GΩ, n = 27 neurons) (Figure 1I-L). There was no significant change of these intrinsic properties of MSNs from DIV 40-75 (RMP: p = 0.265; R_in_: p = 0.09; all were analysed with one-way ANOVA with Bonferroni multiple comparison correction test) with the exception of cell capacitance between DIV 40-45 and DIV 70-75 (p = 0.009). Moreover, we observed firing of action potentials (AP_s_) with the characteristic delayed first spike (Figure 1G) upon current injections. The proportion of functionally mature neurons that displayed evoked APs compared to those that were unresponsive increased as the culture aged (DIV 40-45: 7/10 neurons; DIV 55-60: 20/14; DIV 70-75: 14/6) (Figure 1L). These data indicated robust generation of functionally active iPSC-MSN monocultures that are electrophysiologically stable from DIV 40 to DIV 75.

### In vitro recapitulation of the striatal microcircuit on custom microfluidic devices

Despite the functionally active status of iPSC-derived MSNs in monocultures, we noted a more depolarised RMP and larger R_in_ in *in vitro* MSNs relative to *in vivo* MSNs ^23^. We hypothesised that such discrepancy was principally due to the lack of non-autologous synapses that endogenously provide valuable developmental cues to modulates MSN functional maturation.

To address the lack of maturational signalling within MSN monocultures, we reconstructed the striatal microcircuit *in vitro* mimicking part of the pre-synaptic complexity of the glutamatergic inputs on MSNs using polydimethylsiloxane-based open three-chamber microfluidic chips. The device consisted of three rectangle chambers (4*10 mm) connected via an array of approximately 120 microgrooves of 5 μm wide, 450 μm long and 10 μm high (Figure 2A). The narrow width of the micro-channels compartmentalised the neuronal soma in its respective chamber while DaN and CN axons, but not their dendrites, traversed the full 450 μm length of the microchannels to make connections with MSNs in the middle chamber (Figure 2D).

**Figure 2:**
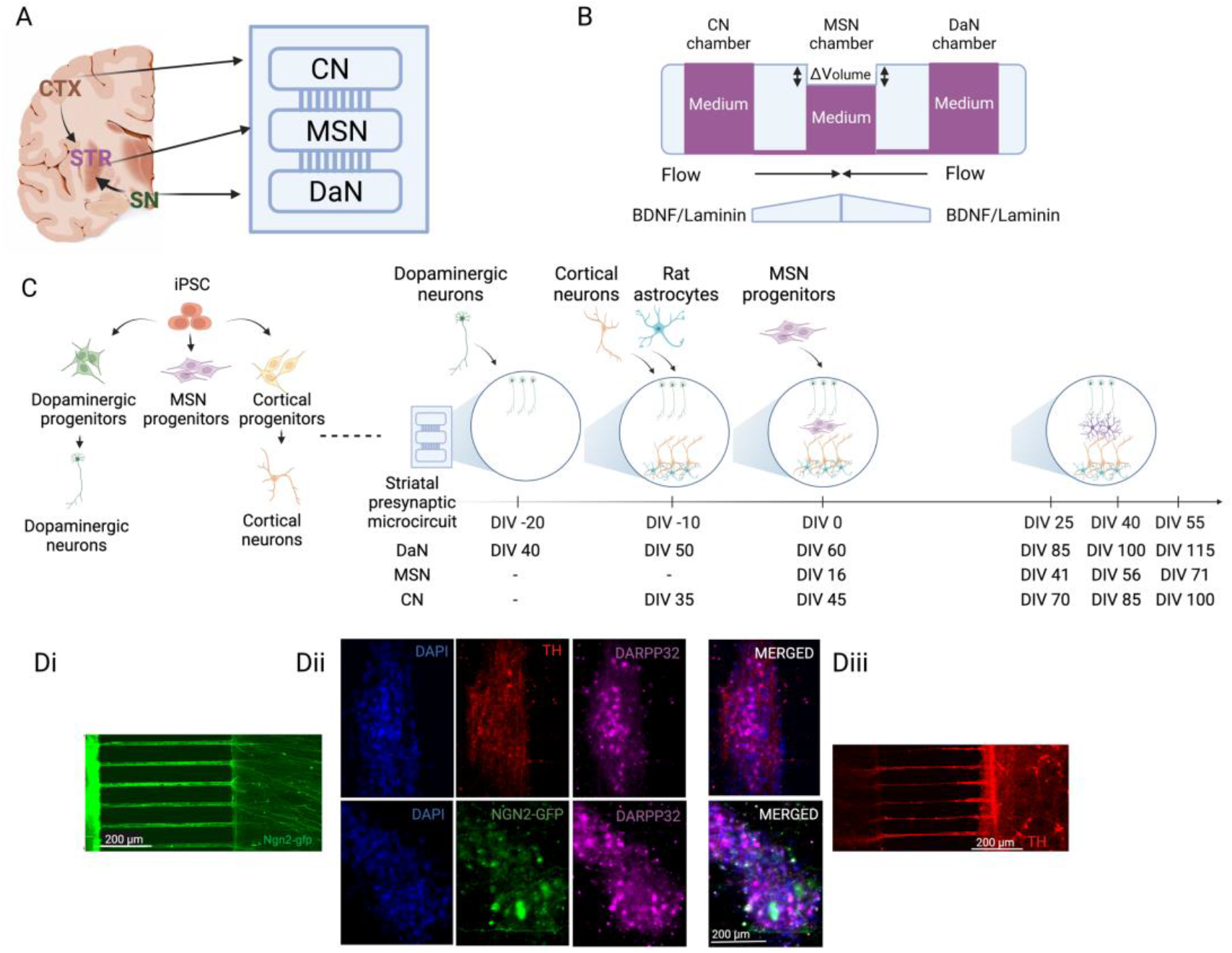
*In vitro* recapitulation of directed striatal presynaptic triad using microfluidic-based devices (A) Schematic of custom microfluidic device which mimics orientation of the cortico-striato-nigral circuit. (B) Side view schematics of fluid and growth factor-driven culturing paradigm of the cortico-striato-nigral microcircuit. (C) Experimental design schematic showing the culturing workflow from iPSC stage to reconstruct the directional connectivity of striatal presynaptic triad. (D) Representative image showing compartmentalisation of (Di) cortical neurons (CNs), (Dii) MSNs and (Diii) dopaminergic neurons (DaNs) in their respective chamber and uni-directional traversing of axons through microchannels.

iPSC-derived DaNs, CNs, MSNs and rat astrocytes were sequentially plated into their respective chambers so as to recapitulate the cortico-striato-nigral triad in the striatum (Figure 2C-D). DaN and CN axons were physically guided toward the middle MSN chamber via flow driven by pressure difference (Figure 2B). We exploited the previous observation that BDNF and laminin are selectively critical for axon polarisation and outgrowth ^24,25^ by setting up artificial BDNF and laminin gradients across the channels to chemically attract axonal projections from the CN and DaN chambers into middle MSN chambers (Figure 2B) prior to MSN deposition. Dopaminergic and cortical axons took approximately 10 days *in vitro* to fully cross and occupy all available microgrooves. Growth of MSN axons travelling out to the DaN and CN chambers was restricted by delaying seeding of iPSC-MSNs until after all microgrooves had been populated with DaNs or CNs axons (Figure 2D). By DIV 0 of the microcircuit culture, the axons of both the iPSC-DaNs and iPSC-CNs had densely projected into the MSN chamber (Figure Di-iii) to regulate MSN progenitor development. In addition to attaining directed connectivity, the culturing schedule was also designed to incorporate known temporal features of these neurons as characterised above in their respective monocultures. Specifically, once the culture platform was established, we applied the strategy of adding DIV 16 iPSC-MSN progenitors to complete the circuit with post-mitotic and functionally active iPSC-DaNs and iPSC-CNs. Cultures were then left until our experimental timepoints (i.e. DIV 40-70 MSNs) when all three neuronal populations were individually electrically active ^26^.

### Glutamatergic input facilitates synaptic maturation of in vitro MSNs

Cortical and dopaminergic afferents form characteristic presynaptic triads with MSN dendrites, specifically on MSN dendritic spine heads and necks, respectively ^27,28^. Corticostriatal terminals represent the major source of glutamatergic input to regulate striatal GABAergic output while dopamine is thought to exert a modulatory influence on the responsiveness of MSNs to excitatory synaptic signals via the regulation of the cell intrinsic excitability state ^23^.

To examine the physiological role cortical collaterals exerted onto iPSC-MSNs in our *in vitro* microcircuit, we compared various electrophysiological parameters between iPSC-MSNs grown on coverslips in monoculture (i.e. CN^−^MSN) versus those co-cultured with iPSC-CNs (i.e. CN^+^MSN) on microfluidic devices (Figure 3A). We found no differences in RMP across time or between culture conditions over the period from DIV 40-70 (RMP: culturing condition p = 0.46; DIV p = 0.29). No significant effect of culturing condition was observed for both cell capacitance and rheobase current at all three timepoints DIV 40-45, 55-60 and 70-75 (CN^−^MSN versus CN^+^MSN: all p > 0.05) (Figure 3B-D). CN^+^MSN also displayed no differences in spontaneous inhibitory post-synaptic current (sIPSCs) (all p > 0.05) (Figure 3E, F and I). However, coculture with CNs exerted significant effect on both frequency and magnitude of spontaneous excitatory post-synaptic currents (sEPSCs), particularly more pronounced in late cultures (sEPSC frequency: DIV 40-45 and 70-75 p > 0.9, DIV 55-60 p = 0.012; sEPSC magnitude: DIV 40-45 p > 0.9, DIV 55-60 p = 0.039, DIV 70-75 p = 0.025) (Figure 3G, H and J). These results imply that our microcircuit setup enables cortical afferents to make functional synapses with MSNs and enhance MSN excitatory synaptic activities.

**Figure 3:**
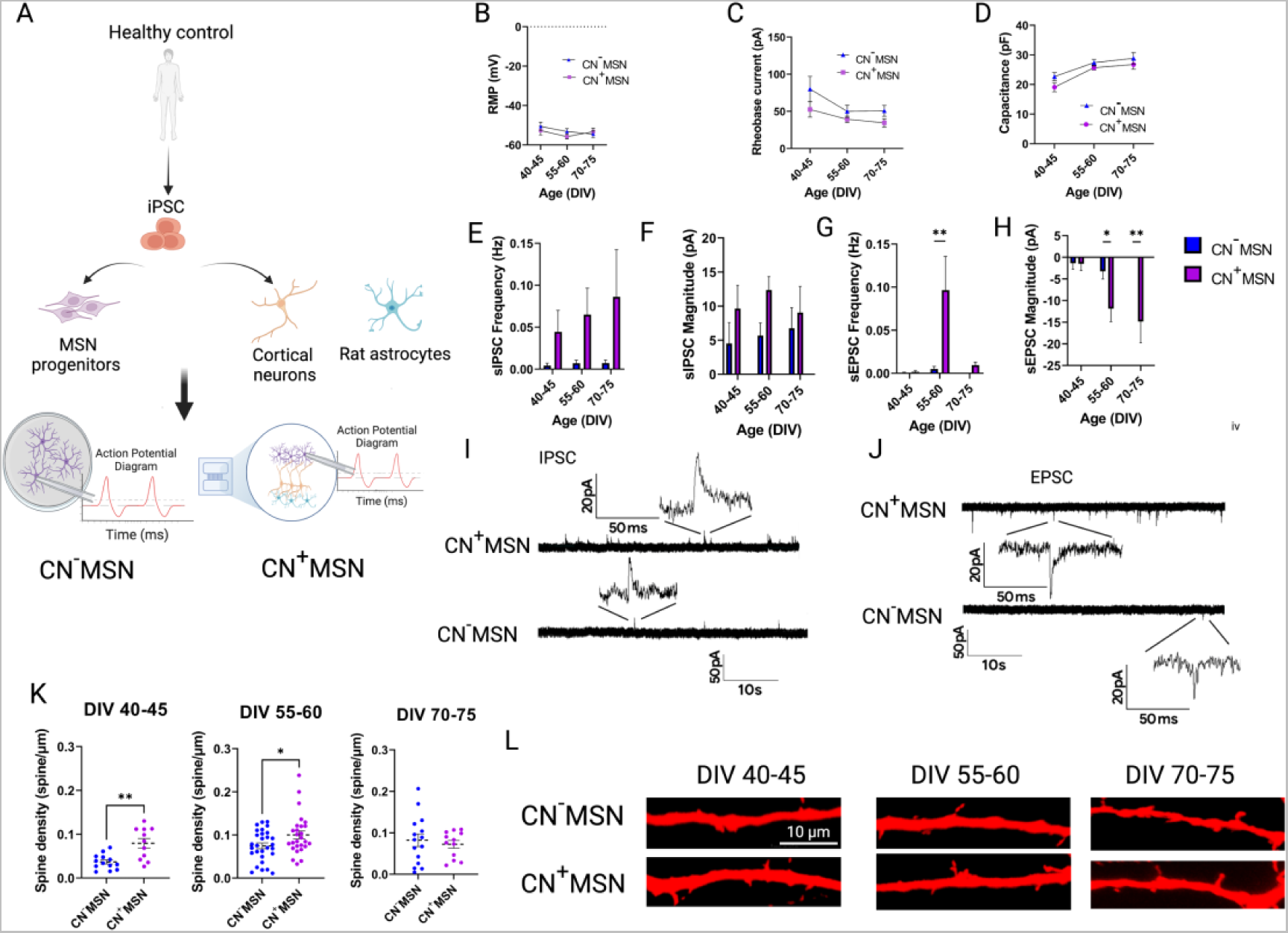
Coculturing MSNs with iPSC-derived cortical neurons improved synaptic activity and dendritic spine morphology of iPSC-derived MSNs. (A) Schematic illustrating the generation and whole cell patch-clamp recording of iPSC-derived MSNs in monoculture (CN^-^MSN) or in co-culture with cortical neurons in microfluidic devices (CN^+^MSN). (B-D) (B) Resting membrane potential (RMP), (C) rheobase current, and D) input resistance of CN^−^MSN and CN^+^MSN, n = 12-42, 7-27, and 12-42 recording MSNs per condition, respectively. (E-H) Quantification of spontaneous inhibitory post-synaptic current (sIPSC) (E) frequency (F) and magnitude over time from DIV 40-75, n = 10-37 recording MSNs per condition. (I) Representative traces of IPSC of CN^−^MSN and CN^+^MSN. (G-H) Quantification of spontaneous excitatory post-synaptic current (sEPSC) (G) frequency and (H) magnitude of CN^−^MSN and CN^+^MSN over time from DIV 40-75. N = 9-21 recording MSNs per condition. (I) Representative trace of EPSC of CN^−^MSN and CN^+^MSN. (K) Pair-wise comparative quantification between CN^−^MSN and CN^+^MSN from DIV 40-75, each dot represents a recording neuron, n = 11-29 recording MSNs per condition, unpaired two-tailed Student’s t-test. (L) Representative images of dendritic spines in neurobiotin-filled DARPP32+ MSNs. All data is presented as mean ± sem, N = 3 differentiation experiments of 3 iPS cell lines, 2-way ANOVA with Bonferroni post-hoc correction test unless otherwise stated. *p<0.05, **p<0.01ßß

Dendritic spines are the principal sites of synaptic integration and their architecture is highly dynamic, reflecting the complex synaptic communication between neurons. Knowing that glutamatergic input profoundly modulates dendritic morphology of MSNs ^29^, we examined whether we could observe a modified phenotype in dendritic spines in the presence of cortical glutamatergic innervation. We found that cortical excitatory afferents promoted early dendritic spine formation in MSNs which was detectable from DIV 40 to 75 (Figure 3K and L). On the contrary, dendritic spines were virtually absent in early culture of MSNs devoid of cortical projections, but emerged gradually over time and attained a similar density as in CN^+^MSN at DIV 70-75 despite CN^−^MSN remaining relatively synaptically more quiescent (DIV 40-45: p = 0.001; DIV 55-60: p = 0.0254, DIV 70-75: p = 0.607 CN^−^MSN vs. CN^+^MSN) (Figure 3K and 2D). These results suggested the importance of cortical afferents in facilitating synaptic maturation of MSNs by prompting not only the formation but also the maturation of dendritic spines of striatal neurons, indicating the utility of our microcircuit in mimicking functional significance of the network connectivity.

### Dopaminergic signalling promoted excitability maturation in a time-sensitive manner

To elucidate the direct effect of dopaminergic projections onto corticostriatal neurons, comparisons were made between CN^+^MSN (i.e. DaN^−^CN^+^MSN) and those circuited with DaNs (i.e. DaN^+^CN^+^MSN, Figure 4A). We found that RMP of corticostriatal neurons became significantly more hyperpolarised with the long-term presence of DaN projections (DIV 40-45 DaN^+^CN^+^MSN vs DIV 70-75 DaN^+^CN^+^MSN p = 0.016), resulting in a significantly more hyperpolarised RMP of DaN^+^CN^+^MSN in late, but not early, cultures (DaN^−^CN^+^MSN RMP vs. DaN^+^CN^+^MSN RMP: DIV 40-45 p = 0.4041, DIV 55-60 p = 0.1120, DIV 70-75 p = 0.044) (Figure 4B). This time-dependent hyperpolarisation of DaN^+^CN^+^MSN RMP was, however, not coupled with differences in rheobase currents and input resistance of MSNs receiving dopaminergic inputs (Rheobase current: culturing condition p = 0.4607; input resistance: culturing condition p = 0.3802) (Figure 4C and D). Observing the trends toward reduced intrinsic excitability produced by dopaminergic afferents, we investigated the constituent voltage-gated currents of sodium (Na_v_) and potassium (K_v_) to delineate specific ionic channels responsible for the subtle differences. The former underlies membrane depolarisation and hence, firing probability ^30^, while the extensive expression of the latter is fundamental for the resting hyperpolarised state of *in vivo* MSNs ^31^. We found an increase in both the fast A-type and slow activating K_v_ in late cultures at DIV 70-75 (DaN^−^CN^+^MSN vs DaN^+^CN^+^MSN fast A-type K_v_ : DIV 40-45 p = 0.508, DIV 55-60 p = 0.317, DIV 70-75 p = 0.0061; DaN^−^CN^+^MSN vs DaN^+^CN^+^MSN fast slow activating K_v_ : DIV 40-45 p = 0.694, DIV 55-60 p = 0.293, DIV 70-75 p = 0.0023) (Figure 4F-H) but no significant temporal changes in Na_v_ currents (DaN^−^ CN^+^MSN vs DaN^+^CN^+^MSN: DIV 40-45 p = 0.927, DIV 55-60 p = 0.794, DIV 70-75 p = 0.235) (Figure 4E), in agreement with the more hyperpolarised RMP but similar rheobase current. The difference in K_v_ can result from either increased K_v_ of DaN^+^CN^+^MSN, or decreased K_v_ of DaN^−^CN^+^MSN, or both. When examining each condition independently over the period of DIV 40-70, we observed a gradual increase in K_v_ among DaN^+^CN^+^MSN while K_v_ of DaN^−^CN^+^MSN increased gradually between DIV 40-65 but subsequently dropped at DIV 70-75 (Supplementary Figure 3). These data suggest that MSN excitability matures in DA-independent (early) and DA-dependent (late) stages of which the latter is likely to be regulated via extensive DA-mediated K_v_ conductances.

**Figure 4:**
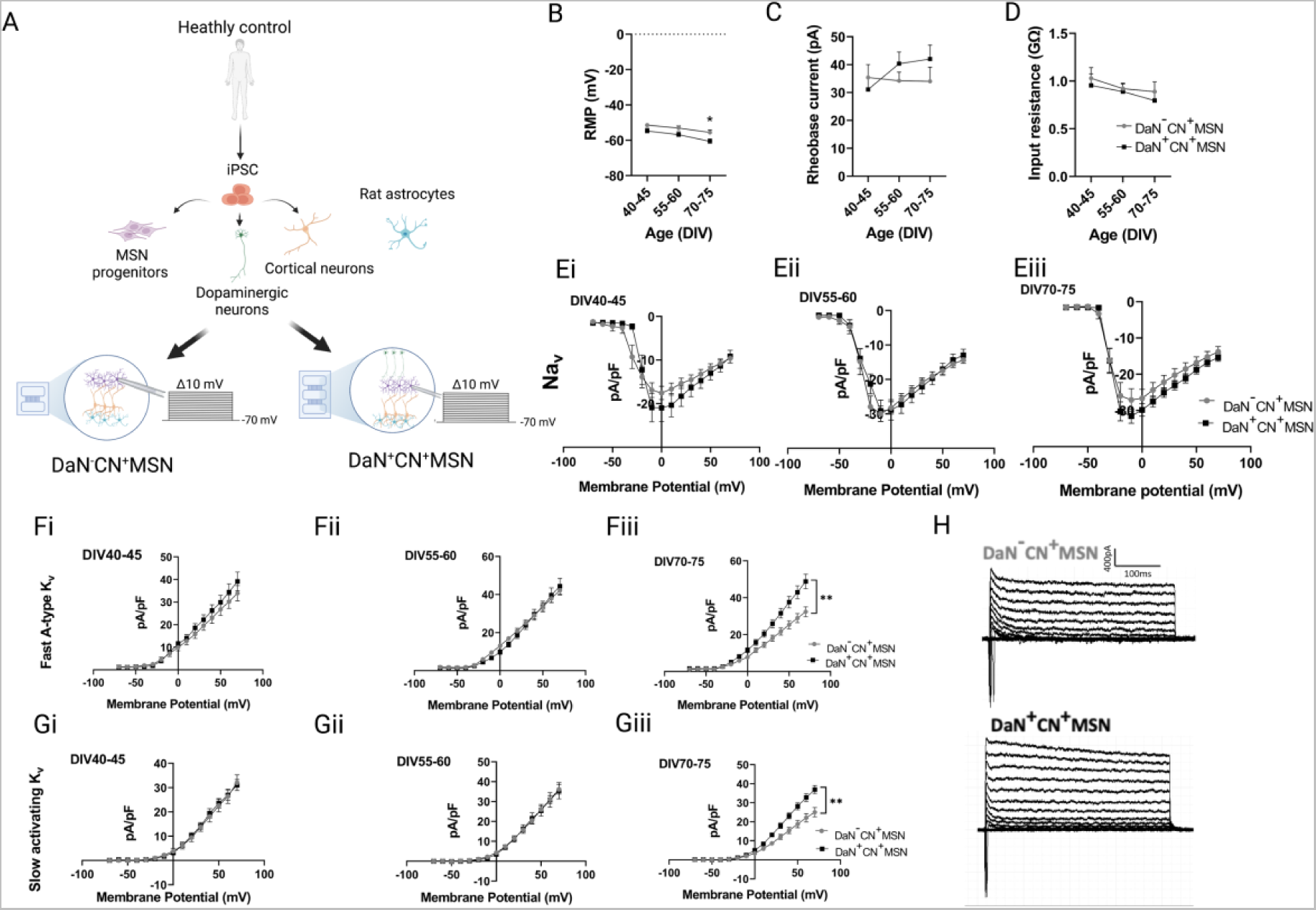
Dopaminergic neurons enhanced the intrinsic maturation of iPSC-derived MSNs during maturation via maintenance of high potassium conductances. (A) Schematic illustrating the generation and whole cell patch-clamp recording of corticostriatal neurons in the presence or absence of DaNs, i.e. DaN^−^CN^+^MSN and DaN^+^CN^+^MSN, respectively, on microfluidic devices. (B-D) (B) Resting membrane potential (RMP), (C) rheobase current, and (D) input resistance over time, n = 9-35 recording MSNs per condition. (E-G) Current density curves of (E) voltage-gated sodium channels, (F) fast A-type voltage-gated potassium, and (G) slow activating voltage-gated potassium channels over time, n = 19-30 recording MSNs per condition. (H) Representative fast A-type and and slow activating K_v_ currents of DaN^−^CN^+^MSN and DaN^+^CN^+^MSN at DIV 70-75. All data is presented as mean ± sem, N = 2 differentiation experiments of 3 iPS cell lines, 2-way ANOVA with Bonferroni post-hoc test, *p<0.05, **p<0.001.

### Early transient electrophysiological deficits of corticostriatal neurons circuited with GBA-N370S DaNs

*GBA1* mutations are a major genetic risk factor for PD with the N370S (the c.1226A > G) variant as one of the most common worldwide ^32^. Multiple previous studies have proposed autonomous cellular dysfunction of DaNs harbouring *GBA-N370S* with a growing literature suggesting failure of the mitochondria and endolysosomal pathway ^7,8,12^. However, it remains elusive as to whether and how the failing mutant DaNs contribute towards physiological perturbation of the striatal output and the eventual motor impairment in PD.

To dissect the impact of *GBA-N370S* iPSC-DaNs on the physiology of corticostriatal output neurons, we established the micro-striatal circuit using iPSC-derived CNs and MSNs from healthy controls connected with iPSC-DaNs from either healthy controls or PD patients carrying the *GBA-N370S* mutation (Figure 5A). We compared the electrophysiological properties between corticostriatal neurons in microcircuit with either healthy DaNs (DaN^+^CN^+^MSN) or *GBA-N370S* DaNs (*GBA-N370S* DaN^+^CN^+^MSN). We found that while both groups exhibited increasingly hyperpolarised RMP over time, the RMP of DaN^+^CN^+^MSN was always more hyperpolarised than that of *GBA-N370S* DaN^+^CN^+^MSN, particularly statistically pronounced in early culture (DIV 40-45: p = 0.0022, DIV 55-60 p = 0.0204, DIV 70-75 p = 0.252) (Figure 5B). Similarly, a significant genotype effect was observed for rheobase current at DIV 40-45, but not in later cultures (DIV 40-45: p = 0.0143, DIV 55-60 p > 0.99, DIV 70-75 p = 0.319) (Figure 5C).

**Figure 5:**
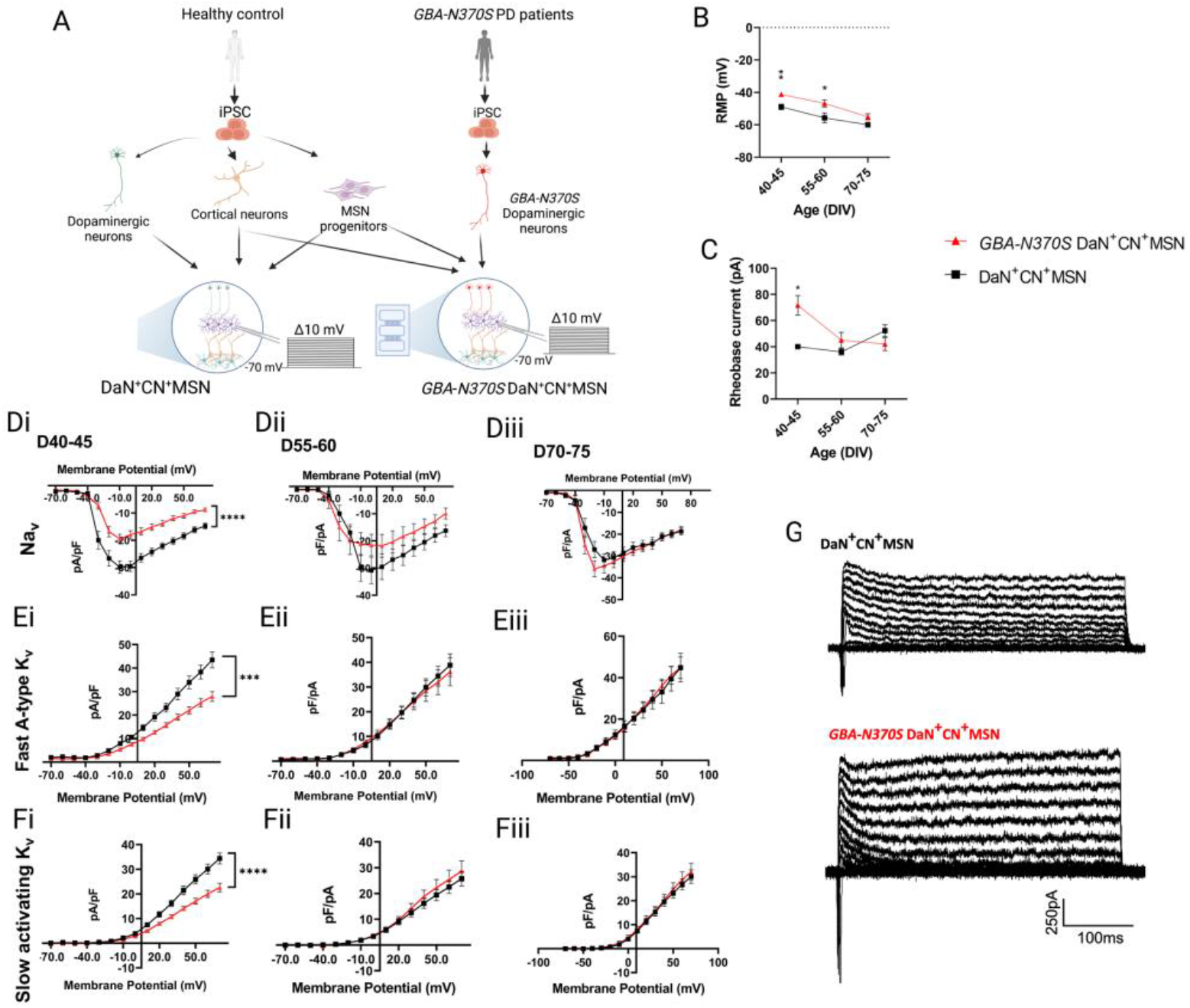
Early but transient current deficits and altered excitability of MSNs circuited with *GBA-N370S* DaNs. (A) Schematic illustrating the generation and whole cell patch-clamp recording of DaN^+^CN^+^MSN and *GBA*-*N370S* DaN^+^CN^+^MSN. (B-C) (B) Resting membrane potential (RMP), and (C) rheobase current of DaN^+^CN^+^MSN and *GBA*-*N370S* DaN^+^CN^+^MSN over time, n = 5-38 recording MSNs per condition. (D-F) Current density curves of (D) voltage-gated sodium channels, (E) fast A-type voltage-gated potassium, and (F) slow activating voltage-gated potassium channels over time. (G) Representative differential fast A-type and and slow activating K_v_ currents of DaN^+^CN^+^MSN and *GBA*-*N370S* DaN^+^CN^+^MSN at DIV 40-45. All data is presented as mean ± sem, N= 2-3 differentiation experiments of 3 iPS cell lines, n = 27-32 recording MSNs at DIV 40-45, n = 14-16 recording MSNs per condition at DIV 55-60 and DIV 70-75. 2-way ANOVA with Bonferroni post-hoc test, ns p>0.05, *p<0.05, **p<0.01, ***p<0.001, ****p<0.0001.

Neuronal excitability is tightly controlled by coordinated activity of both Na_v_ and K_v._ The inward flux of sodium ions depolarises the cell membrane towards action potential threshold and triggers spontaneous firing while outward currents generated K_v_ underly resting membrane potentials and action potential afterhyperpolarisation ^30^. Hence, we next explored whether the altered intrinsic excitability was coupled with relevant ionic channels. Unsurprisingly, we observed a substantial reduction in current density of both Na_v_ and K_v_ among the *GBA-N370S* DaN^+^CN^+^MSN (DIV 40-45: Na_v_ and slow activating K_v,_ DaN genotype p <0.0001; fast A-type K_v_, DaN genotype p = 0.0001) (Figure 5Di, Ei, Fi, and G). Recordings of iPSC-MSNs from intermediate and late cultures between DIV 55-70 revealed resolution of genotypic effects previously observed at DIV 40-45 (Figure 5D-F). These data suggest an early pathological role of *GBA-N370S* mutation in iPSC-DaNs in altering striatal GABAergic output via early hyperexcitability of post-synaptic iPSC-MSNs independent of overt degeneration of dopaminergic afferents.

### PKA antagonism resolved early and transient ionic current deficits in corticostriatal neurons circuited with GBA-N370S iPSC-DaNs

The electrophysiological activity of many cellular channels, including voltage-gated sodium and potassium channels, is robustly regulated by PKA-mediated signalling pathway in response to fluctuations of external stimuli ^33,34^. In MSNs both K_v_4.2 and K_v_1.2, the principal channel contributing towards fast A-type and slowly inactivating potassium current, respectively, as well as Na_v_, are negatively modulated by PKA activation ^35–38^ To examine whether PKA activity is involved in the early deficits of corticostriatal neurons affected by *GBA-N370S* iPSC-DaNs, we applied the cell-permeable PKA inhibitor H89 directly to iPSC-MSNs in the striatal chambers of the striatal microcircuit and performed whole-cell patch clamping of iPSC-MSNs at DIV 40-45 (Figure 6A and B). PKA activity assay data suggested an intrinsic doubling of PKA activity in iPSC-MSNs circuited with *GBA-N370S* iPSC-DaNs (i.e. *GBA-N370S* DaN^+^CN^+^MSN), as compared to those with healthy control-derived iPSC-DaNs (i.e. DaN^+^CN^+^MSN) (DaN^+^CN^+^MSN vs *GBA-N370S* DaN^+^CN^+^MSN p = 0.0426) (Figure 6C). The increase in PKA activity in the iPSC-MSNs was then reduced to the control level upon administration of 10 μM H89 to iPSC-MSNs for 15 mins (DaN^+^CN^+^MSN vs *GBA-N370S* DaN^+^CN^+^MSN + 10 μM H89 p > 0.99, *GBA-N370S* DaN^+^CN^+^MSN vs *GBA-N370S* DaN^+^CN^+^MSN + 10 μM H89 p = 0.05) (Figure 6C). These data demonstrated the phenotypic increase in PKA activity in MSNs synapsed with *GBA-N370S* DaNs and the ability of H89 to normalise striatal PKA activity *in vitro*. DaN lysates were collected from the DaN chambers from the same devices as the healthy control-derived striatal lysates, and a significant reduction in PKA activity was seen in *GBA-N370S* DaNs (i.e. MSN^+^CN^+^*GBA-N370S* DaN) relative to healthy control-derived DaNs (i.e. MSN^+^CN^+^DaN) (MSN^+^CN^+^DaN vs MSN^+^CN^+^*GBA-N370S* DaN p = 0.0179) (Figure 6D). This deficit in PKA activity in *GBA-N370S* DaNs remained in the mutant *GBA-N370S* DaNs synapsed with iPSC-MSNs treated with H89 (p = 0.0373 relative to MSN^+^CN^+^DaN), confirming the fluidic isolation of different neuronal compartments. Treatment of iPSC-MSNs with H89 resolved the genetic effect of *GBA-N370S* DaNs on MSNs Na_v_ and K_v_ current density at DIV 40-45 to healthy control levels (Figure 6E-G), indicating the involvement of PKA in mediating intrinsic excitability of corticostriatal neurons via ionic conductances.

**Figure 6:**
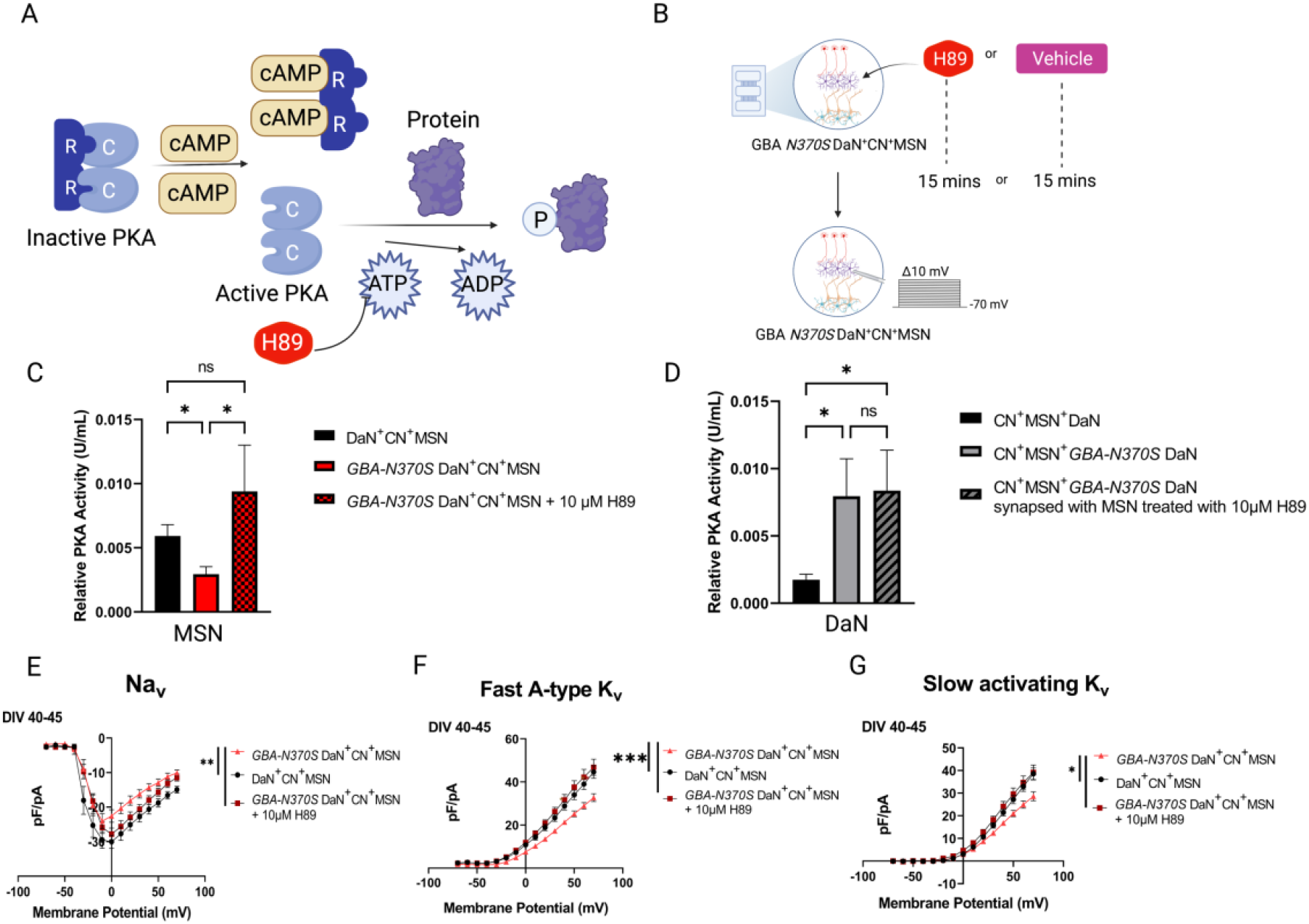
PKA antagonist rescued early current deficits of MSNs circuited with *GBA-N370S* DaNs. (A) Schematic demonstrating antagonistic mechanism of H89 in PKA-mediated phosphorylation signalling cascade. (B) Schematic illustrating treatment of PKA antagonist H89 on MSNs in the striatal presynaptic microcircuit; C = catalytic subunit, R = regulatory subunit, cAMP = Cyclic adenosine monophosphate, ATP = Adenosine triphosphate, ADP = Adenosine diphosphate. (C-D) Quantitative measure of PKA activity of (C) DaN^+^CN^+^MSN, *GBA-N370S* DaN^+^CN^+^MSN and *GBA-N370S* DaN^+^CN^+^MSN treated with PKA antagonist H89, Kruskal-Wallis test with Dunn’s correction test, and (D) healthy DaNs, mutant *GBA-N370S* DaNs synapsed with healthy untreated MSNs or mutant *GBA-N370S* DaNs synapsed with H89-treated MSNs in their corresponding devices; Friedman test with Dunn’s correction test. N = 2 differentiations of 3 iPS cell lines each, n = 3 technical replicates; values were measured by PKA colorimetric activity kit and normalised against total protein level extrapolated from BSA assays. Each unit per ml (U/ml) was defined as the amount of PKA needed to mediate the transfer of 1.0pmol phosphate from ATP to substrate at 30°C as per manufacturer’s instructions. (E-G) Current density curves of (E) Na_v_, (F) fast A-type K_v_, and (G) slowly activating K_v_ of DIV 40-45 MSNs. All data were presented as mean ± sem, N = 2 differentiation experiments of 3 iPS cell lines, n = 17-22 recording MSNs per condition. 2-way ANOVA with Bonferroni post-hoc test unless otherwise stated, ns p>0.05, *p<0.05, **p<0.01, ***p<0.001

## Discussion

The present study represents the first attempt at recapitulating an *in vitro* neuronal circuit model of the human striatal synaptic triad using an open chamber-based microfluidic platform to reveal the physiological importance of connectivity-driven inputs on human striatal functions. Our data suggest an early nonautonomous striatal electrical dysfunction in MSNs in the presence of the *GBA-N370S* mutation within presynaptic DaNs that was associated with diminished striatal PKA activity. Interestingly, in this model striatal neurons showed dynamic cell-autonomous PKA-dependent modulation of ionic channel physiology as illustrated by the resolution of early striatal deficits by specific antagonism of striatal PKA activity, despite the persistent presence of mutant DaNs.

Previous strategies to investigate human striatal neuron (patho)physiology remain largely limited in monocultures of a single neuronal type ^20^, with more recent success in developing a neuronal network architecture ^16^. To date, there are no cellular models that capture the complex triad of non-autologous synapses onto striatal neurons which are critical for the emergence of functional characteristics of MSNs. Indeed, although our MSNs in monoculture displayed a temporal marker expression profile reminiscent of the developmental trajectory reported from the Human Brain Transcriptome (https://hbatlas.org), combined cortical glutamatergic and dopaminergic inputs further facilitated functional maturation of the MSNs beyond the benefits of long-term culturing. Our results corroborated the relevance of cortical glutamate in promoting both inhibitory and excitatory synaptic maturity of MSNs ^39^ as well as the time-sensitive influence of dopaminergic transmission on MSNs excitability maturation previously reported in rodent models ^23^. It should also be noted that the glutamatergic input onto MSNs originates from not only the cortex but also the thalamus which makes up ~40% of glutamatergic synapses on MSN dendrites and converges with corticostriatal afferents indiscriminately onto both MSN subtypes ^40^.

A significant highlight of this open microfluidic-based circuit approach is its customisability. The modular nature of this setup allows great flexibility for reconstructing the compartmentalisation of different neuronal populations and the directionality of their specific connectivity. Hence, virtually any microcircuit can be established applying a similar design concept. Targeted manipulation to specific compartments is also permitted with great ease as exemplified by our striatal PKA treatment paradigm. Immunostaining and electrophysiological results indicate that our approach has captured the oriented connectivity resembling that *in vivo* and that functional glutamate and dopamine transmission exists.

Because the ATP competitive analogue H89 produced similar amplification of both Na_v_ and K_v_ in our study, the observed elevation of striatal PKA activity likely occurred upstream of the cAMP in the PKA signalling cascade, potentially through excessive Dopamine 1 receptor (D1R) activation followed by stimulation of adenylyl cyclase-cAMP cascade ^41^. The D1R antagonist SKF83566 has been shown previously to decrease PKA activity in direct pathway MSNs ^42^. Voltage clamp work has confirmed that D1R stimulation reduced Na_v_ currents by limiting channel availability ^43^ while lessening K_v_ peak current amplitude through altering channel inactivation kinetics ^38^. Furthermore, the D1R agonist SFK81297 reversibly reproduced the reduction in Na_v_ and K_v_ current amplitude, similarly through PKA-mediated phosphorylation of channel subunits, and improved striatal firing probability ^44,45^.

The majority of our culture was composed of direct pathway MSNs (dMSNs) expressing mainly substance P or pro-dynorphin (Supplementary Figure 2) implicating a potential role for D1R. Considering that the striatal excitability in the absence of dopaminergic projections at DIV 40-45 (Figure 4Ei, Fi, and Gi) did not phenocopy the early striatal deficit synapsed with *GBA-N370S* DaNs at the same time point (Figure 5Di, Ei, and Fi), we speculate that increased, rather than reduced, tonic DA content at the nigrostriatal synapses is a more logical explanation of D1R over-stimulation. This hypothesis aligns with previous work on other models of PD-related familial mutations reporting early cell-autonomous hyperexcitability of midbrain DaNs prior to overt neurodegeneration ^46–48^ which could result in increased PKA activity in dMSNs ^42^, but contrasts with a single report on *GBA-N370S*-associated reduced spontaneous firing and basal DA release by Woodard and colleagues ^49^ in monocultured DaNs.

Epidemiological data suggests that only about 9.1% of *GBA1* mutation carriers will develop PD, often with a relatively earlier age of onset and modest motor impairment as compared to non-carriers ^32^. This is consistent with the early changes in striatal function observed in our study which was subsequently resolved at later timepoints. As the compensation was likely to be intrinsic to non-mutant MSNs, we propose that *GBA1* mutation-driven deficits in DaNs emerge early but in themselves are insufficient to induce long-term dysfunction in the motor circuit regulated by the basal ganglia. This evidence of potential compensative capacity is in keeping with the late age of disease presentation in PD. It is interesting to speculate that MSNs might be able to compensate for PD disease-driving deficits elicited by DaNs, thereby masking initial disease stages and the prodrome. A functional motor study of reversible PKA modulation *in vivo*, would be an interesting experiment to further test these compensation effects in a PD mouse model that was not yet symptomatic for motor impairments. While major therapeutic efforts have been channelled toward reversing DaN pathology, our results suggest that rescue of the striatal deficits is an attractive alternative to preserve the integrity of the basal ganglia output. If compensation is indeed functionally implicated in genetic disease as our data suggests, then therapies targeting MSNs might be able to provide patients with symptomatic relief. Future studies could mimic the full genetic landscape of *GBA-N370S* carriers by deriving all three neuronal populations of the microcircuit from the same *GBA-N370S* cell lines. Pairwise comparisons of striatal function from microcircuits in which the *GBA1* mutation burden is carried by different, all, or none of the component cell types mixed with healthy controls could delineate contributing pathophysiological roles of different neuronal populations of the striatal presynaptic circuitry to the net debilitating burden of the *GBA1* mutation.

We have demonstrated a PKA activity-related mechanism by which mutant *GBA-N370S* DaNs induced early hyperexcitability of healthy control-derived iPSC-MSNs preceding neurodegeneration. We also observed a reduction in PKA activity in *GBA-N370S* iPSC-DaNs relative to healthy control-derived DaNs (Figure 6D) which is consistent with the previously reported downregulation of PKRACB transcript expression from DaN monocultures harbouring the same genetic burden ^8^. Mutation of the PKA regulatory subunit *PRKAR1B* has also been reported in a novel familial neurodegenerative condition with parkinsonism and dementia ^50^, and *Prkar1b*-knockout rats exhibit body tremor ^51^. Moreover, reduced PKA activity in DaNs has been implicated in PD pathophysiology in other PD-related rodent mutation models such as *LRRK2* R1444C/G/H ^52,53^ and *PINK1* ^54^. Likewise, increased PKA activity was thought to be protective against PD ^42,55^. However, the exact mechanistic link between *GBA1* mutations and PKA activity in nigral neurons remains to be further investigated. Nevertheless, it is well documented that the large network of lysosome genes including *GBA1* is moderated by a master regulator transcription factor EB (TFEB) ^56^ which in turn is the downstream substrate of PKA-mediated phosphorylation via mTORC1 ^57^. Considering our previous observation indicating dysregulation of lysosomal/autophagic signalling in monocultured DaNs carrying *GBA-N370S* ^7,8^, we suggest that the multiple cellular perturbations in nigral DaNs burdened by the *GBA-N370S* mutation might be initiated by reduced activity of the master kinase PKA which disrupts various associated neuronal functions potentially via lysosomes and/or lysosomal-related organelles. In addition, PKA activation is closely related to proteasomal activation ^58,59^, which is a necessary function often compromised in PD ^60,61^. It is also interesting to speculate that reduced PKA activity could alter protein homeostasis through reduced proteasomal function, and so, in turn, cause cellular dysfunction. From a preferential vulnerability point of view, protected cells like MSNs might be able to find compensation mechanisms and recover from these effects, while vulnerable cells such as DaNs may not be able to ^62^.

PKA was thought to positively modulate neurotransmitter release in most synapses via a combination of factors including phosphorylation of various proteins involved in synaptic vesicle exocytosis ^63–65^. *In vivo* studies have suggested PKA phosphorylation-mediated increase in DA release via activation of TH enzymatic activity ^66,67^. The involvement of PKA in neurotransmitter release suggests that mutant *GBA-N370S* may affect DA striatal content, and that the pathophysiology of *GBA-N370S* variant-mediated PD may result from a disruption of the tight modulation of PKA activity.

In conclusion, we have described an open microfluidic-based neuronal microcircuit model which has used iPSC-derived neurons to recapitulate *in vitro* the presynaptic network relevant to *in vivo* striatal neurons. This approach allowed us to explore emerging electrophysiological features of human MSNs over time in both the healthy and diseased state. We envision greater use of this modular circuit-based model in addressing sophisticated functional questions of human physiology and pathology.

## Method details

### 1. Acquisition and maintenance of iPS cell lines

iPS cell lines used in this study were derived from human skin biopsy fibroblast acquired with informed consent and ethical approval (Ethics committee: National Health Service, Health Research Authority, NRES Committee South Central, Berkshire, UK, REC 10/H0505/71). The iPS cell lines have been previously characterized: 3 control iPSC lines SFC067-03-01 (RRID:CVCL_RD75), SFC156-03-01 (EBiSC Cat# STBCi101-A, RRID:CVCL_RD71) ^8^, and SFC856-03-04 (RRID:CVCL_RC81), 3 *GBA*-*N370S* PD lines MK082-26 (RRID:CVCL_IJ04) ^8^; MK088-01 (EBiSC Cat# UOXFi003-A), and MK071-03 (EBiSC Cat# UOXFi001-B) ^7^. All iPSCs were maintained on Matrigel™ (Corning)-coated plates under feeder-free culture condition with mTSER1™ (StemCell Technology) supplemented with 1% penicillin/streptomycin (P/S) (Life Technologies). Medium was changed daily, and cultures were passaged 1:2 every other day using TrypLE Express™ (Life Technologies) when they reached 90-100% confluency in the presence of Rock inhibitor (Bio-Techne).

### 2. Generation of iPSC-derived MSNs

3 iPSC control lines (SFC067-03-01, SFC156-03-01, and SFC856-03-04) were differentiated into MSNs using condition modified from previously established protocols ^14,15^. In brief, iPSCs were plated at 75,000 cells/cm^2^ and maintained until 85% confluence. Culture was switched to neural induction medium containing DMEM/F12 (Life Technologies), 1% MEM Non-Essential Amino Acids (NEAA), 1% Glutamax, 2% B27 without vitamin A (all from ThermoFisher), 1% P/S, 1 μM LDN193189 (Sigma), 10μM SB431542 (Abcam) and 4 μM XAV (Tocris). On DIV 04, culture was pre-incubated with ROCK Inhibitor (Y-27632) for 1 hour before being passaged in 1:2 ratio. Culture was passaged in 1:2 on DIV 8 and switched to differentiation medium containing DMEM/F12, 1% P/S, 1% NEAA, 2 mM L-glutamine (Life Technologies), 2% B27, 200 nM LDN193189, and 4 μM XAV. Activin A was added to from DIV 12 till DIV 23.

On day 16, the culture was either cryopreserved for future experiments or replated at 65,000 cells/cm^2^ onto freshly coated plates or coverslips. Coating included 10 and 100 μg/ml Poly-D-Lysine (Sigma-Aldrich) overnight for plastic plates and glass coverslips, respectively, followed by 1:100 Matrigel™ for 1 hour. Neuronal progenitor cells were cultured in the maturation medium SynaptoJuice® as previously reported ^15^. Cultures were treated with 1 μM AraC on DIV 18 to arrest proliferation of non-neuronal cells capable of DNA synthesis. AraC concentration in the culture was gradually diluted with subsequent media changes.

### 3. Generation of iPSC-derived CNs

Generation of CNs from 3 iPSC control lines ((SFC067-03-01, SFC156-03-0, and SFC856-03-04) was adapted from a previously detailed protocol ^26^. In brief, neural development was induced with dual SMAD inhibitors 10 μM SB431542 (Tocris) and 118 nM LDN (Sigma). Following formation of a dense neuroepithelial sheet, the whole sheet was enzymatically lifted using Dispase (Life Technologies) and broken down into smaller clumps to encourage development of neural rosettes. Rosettes were maintained and expanded on laminin-coated wells till DIV 30 at which point cultures were either cryopreserved or dissociated into single cells for further differentiation. At DIV 30 cortical neurogenesis was induced by doxycyline-dependent overexpression of *NGN2* via co-transduction of LV-TetO-mNgn2-T2A-Puro and LV-Ubiq-rTA lentiviruses into the cultures. Transduced cells were incubated for 2 days before being selected with fresh media supplemented with 1 μg/ml puromycin for another 48 hours prior to final replating.

### 4. Generation of iPSC-derived DaNs

All 3 control and 3 *GBA N370S* PD iPS cell lines were differentiated into DaNs as previously described ^8^ with added modifications ^68^. In brief, iPSCs were dissociated into single cells, plated on Geltrex-coated plastic plates at 125,000 cells/cm^2^ and cultured till confluence. iPSCs were then patterned toward floor-ventral midbrain precursors for 10 days. Day 10 precursors were expanded with weekly passaging for the subsequent 20 days. These cells can be used immediately or cryopreserved. Post-expansion DIV 29 precursors were further differentiated and matured into post-mitotic midbrain neurons until DIV 38 for final replating onto microfluidic devices.

### 5. Immunocytochemistry

Cultures were fixed with 2% PFA for 20 mins at room temperature (RT), rinsed with PBS, and antigen retrieved in citrate buffer pH 6.0 (ThermoFisher) in water bath at 80°C for 5 mins. Samples were left to rest for 10 mins at room temperature before being permeabilised and blocked in PBS, 10% donkey serum and 0.01% of Triton X-100 for 10 mins. Incubation of primary antibody in PBS and 10% donkey serum was performed overnight at 4°C. Samples were washed with PBS and subsequently incubated in species-appropriate Alexa Fluor© secondary antibody in PBS with 10% donkey serum for 1 hour at RT. The list of primary and secondary antibodies used for immunostaining was tabulated in supplementary table 2 and 3, respectively. Images were acquired on the Opera Phenix High-content Screening system (PerkinElmer) or the Invitrogen EVOS ™ FL Auto (ThermoFisher) cell imaging system and subsequently processed on Harmony (Perkin Elmer; RRID:SCR_018809) or ImageJ (RRID:SCR_003070), respectively.

### 6. Reverse transcription-qualitative polymerase chain reaction (RT-qPCR)

Total RNA of MSN cultures over time was extracted and purified from cell lysates using RNeasy Mini kit (Qiagen) according to manufacturer’s instructions. Extracted RNA was quantified and assessed on a Nanodrop (DS-11, Denovix). RNA was reverse transcribed into cDNA using superscript III reverse transcriptase kit (Life Technologies). qPCR was performed with standard SYBR green (ThermoFisher) in StepOnePlus thermal cycler (Life Technologies). Full list of primers used are tabled in Supplementary Table 1.

### 7. Microfluidic fabrication

A custom master mould containing both duo- and trio-chambers was manufactured by Microliquid (Spain). The microfluidic device was made from the mould by pouring a silicon elastomer SylGARD 184, Dow Corning © pre-mixed with its curing agent in the 1:10 ratio. Polymerisation occurred when incubating the PDMS-filled mould for 3 hours at 60°C. Final devices were manually cut out from the negative cast and washed with 100% ethanol. Cut-out devices were air-dried in tissue culture hood and subsequently placed on ethanol –sterilised 19 mm-diameter coverslips to form an instantaneously tight seal. Both the chambers and microchannel area were coated with poly-D-lysin (0.1 mg/ml) overnight followed by Geltrex™ (ThermoFisher) prior to cell culturing.

### 8. Electrophysiology

Whole-cell recordings of MSNs on coverslips were done at 27°C with periodic change of external solution following every new recording cell. The external solution contained: 2.4 mM KCl, 167 mM NaCl, 10 mM Glucose, 10 mM HEPES, 1 mM MgCl_2,_ and 2 mM CaCl_2_ adjusted to 300 mOsmol/l and pH 7.36 with NaOH. Neurons were patched with a borosilicate glass pipette (8-12 MΩ) pulled using a Sutter P-97 Flaming Brown puller (Sutter Instrument Company) and filled with an internal solution consisting of: 140 mM C_6_H_11_KO_7,_ 6 mM NaCl, 1 mM EGTA, 4 mM MgATP, 0.4 mM Na_3_GTP, 10 mM HEPES, and 0.01% Neurobiotin (SP-1120) adjusted to 290 mOsmol/l and pH 7.3 using KOH. Data were acquired with MultiClamp 700B amplifier (Molecular Devices) and Digidata 1550B digitiser (Molecular Devices) and analysed in Clampfit 10.7 (Molecular Devices). Cells were always clamped at −70 mV unless otherwise stated. Access resistance was constantly monitored; cells with access resistance greater than 50 MΩ were excluded. All recordings are made within two hours of coverslip preparation. Cell capacitance and input resistance were automatically reported from Clampex membrane test sampled at 33 kHz. Resting membrane potentials (RMP) were acquired immediately upon successful break-in in current clamp with zero current injection. Liquid junction potential was estimated at 12mV using JPCal ^69^ and adjusted for in RMP recordings.

Na_V_ and K_V_ currents were recorded in voltage clamp using a series of 400 ms square voltage steps of 10mV increments from −70 mV to +70 mV. Signals were sampled at 10 kHz and filtered at 2 kHz. Leak subtraction was applied with 4 sub-sweeps and a settling time of 250 ms. Evoked action potentials were recorded in current clamp by injecting 500 ms current steps in 10 pA increments from −10 pA to +130 pA.

Spontaneous EPSC and IPSC were recorded in voltage-clamp mode with 20x gain at one another’s reversal potential i.e. −70 mV and 0 mV, respectively. The internal solution for spontaneous synaptic activity recordings was Cs-based containing 140 mM C_6_H_11_CsO_7,_ 6 mM NaCl, 1 mM EGTA, 4 mM MgATP, 0.4 mM Na_3_GTP, 10 mM HEPES, and 0.01% Neurobiotin adjusted to 290 mOsmol/l and pH 7.3 using CsOH. All sEPSC and IPSC recordings were 1-4 mins in duration and low-pass Bessel 8-pole filtered post-hoc. Spontaneous post-synaptic events and event magnitude were automatically detected based on absolute magnitude difference from the baseline (>10 pA for sEPSCs and >15 pA for sIPSCs) using threshold search function on Clampfit (RRID:SCR_011323). Putative EPSC or IPSC events were excluded based on template-dependent criteria including rise time and half-width (<4 ms and <1.2 ms, respectively) and manually validated to reject false-positive events.

H89 (Tocris) was dissolved in dH2O and were media applied to the middle chamber to the final concentrations of 10 μM for 15 mins prior to wash off and recording.

All patched neurons were post-hoc labelled with Streptavidin-Alexa Fluor 488 conjugate (ThermoFisher) and DARPP32 (Sigma) to ascertain the MSN identity. Only neurobiotinylated neurons that co-expressed DARPP32 were selected for analysis.

### 9. Dendritic spine visualisation and quantification

Patched coverslip of MSNs was immunolabelled for DARPP32 (as described above) and mounted onto SlowFade ™ Diamond Antifade mountant (ThermoFisher). Detection and imaging of DARPP32-positive Neurobiotin™-filled neurons were captured under Olympus FluoView FV1000 confocal microscope with argon and solid-state laser with 488 nm and 559 nm excitation, respectively. Z-stacks of images were sampled sequentially at resolution of 1024 * 1024 pixels, with 60x oil-immersion objective (NA = 1.40), at 1.05 μm steps as optimised by Nyquist sampling theorem. Dendritic branches of biotinylated neurons were captured at 3x software zoom. Images were segmented, reconstructed and automatically rendered using filament and spine detection module in Imaris 9.6.0 (Bitplane, South Windsor, CT, USA, RRID:SCR_007370) which identified dendrites and detect protrusions along the dendritic filament length. Putative spines at branch points or disconnected dots were manually excluded.

### 10. PKA activity assay

PKA activity were measured using the PKA Colorimetric Acitivity Kit (ThermoFisher) according to manufacturer’s guidelines. Protein concentrations of the same samples were quantified using Pierce™BCA Protein Assay Kit (ThermoFisher) following manufacturer’s instructions. Both plates were read for absorbance in a PHERAstar® microplate reader (BMG Labtech). Protein concentration and PKA activity values were inferred from their respective standard curves. Final PKA activity read-outs were normalised against the sample protein concentrations.

### 11. Statistical analysis

All data was presented as mean ± Standard error of the mean (sem) unless otherwise stated. Raw data was tested for normality and statistical comparison of the means was performed using unpaired Student’s t-test, two-way ANOVA with Bonferroni post-hoc test, or Kruskal-Wallis test or Friedman test corrected with Dunn’s multiple comparison test when appropriate. Significance difference was considered at *p* < 0.05. All statistical analyses were performed on GraphPad Prism 6.0 (GraphPad Software, RRID:SCR_002798).

---

**Supplementary Table 1:**
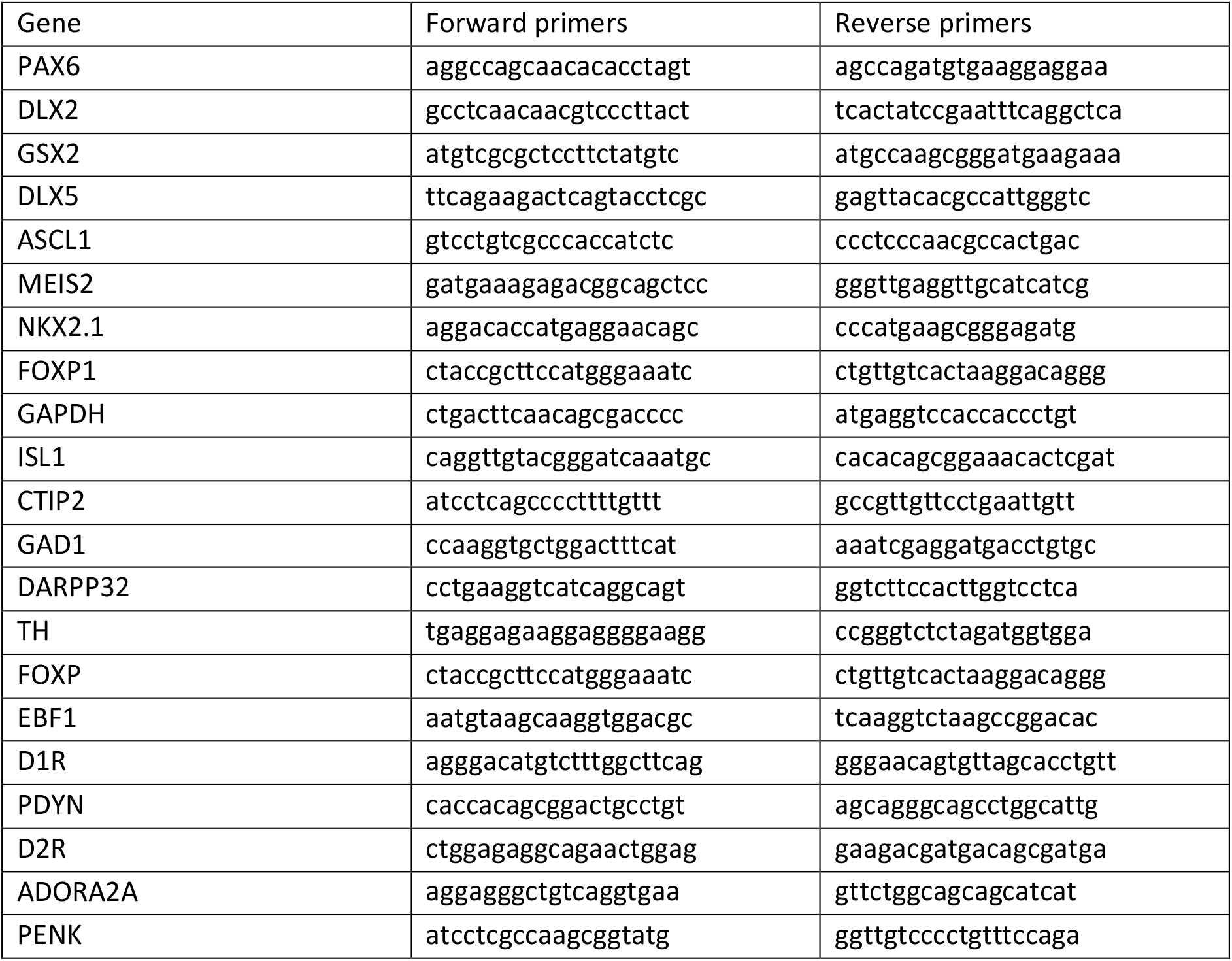
Primer sequences

**Supplementary table 2:**
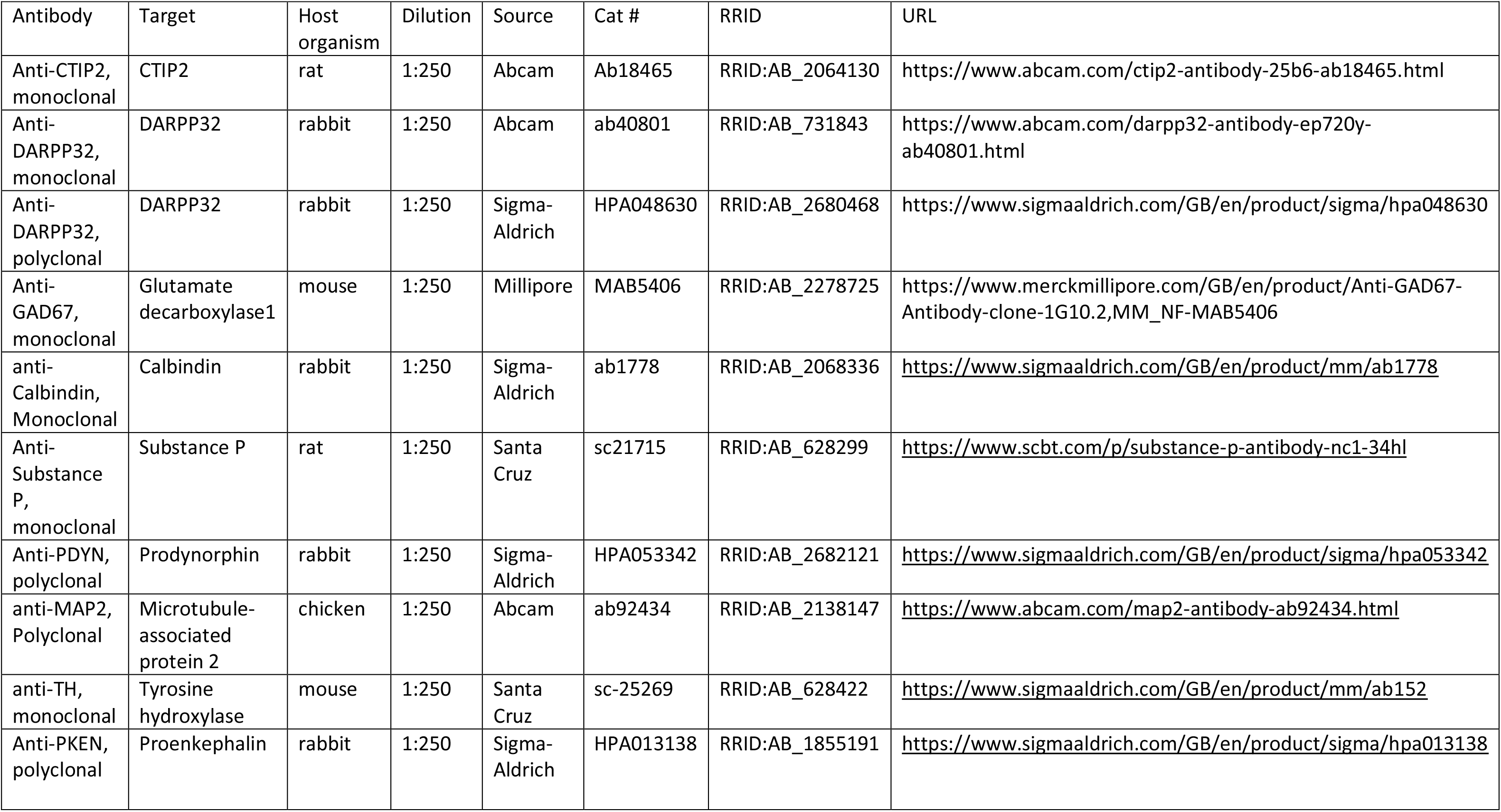
Primary antibodies used

**Supplementary table 3:** Secondary antibodies used.

| Antibody | Target | Host organism | Dilution | Source | Cat # | RRID | URL |
| --- | --- | --- | --- | --- | --- | --- | --- |
| Alexa-Fluor 555 | IgG Mouse | Donkey | 1:1000 | Invitrogen | A-11057 | RRID:AB_2534104 | https://www.thermofisher.com/antibody/product/Donkey-anti-Goat-IgG-H-L-Cross-Adsorbed-Secondary-Antibody-Polyclonal/A-11057 |
| Alexa-Fluor 555 | IgG Rabbit | Donkey | 1:1000 | Invitrogen | A-31570 | RRID:AB_2536180 | https://www.thermofisher.com/antibody/product/Donkey-anti-Mouse-IgG-H-L-Highly-Cross-Adsorbed-Secondary-Antibody-Polyclonal/A-31570 |
| Alexa-Fluor 648 | IgG Rabbit | Donkey | 1:1000 | Invitrogen | A-31572 | RRID:AB_162543 | https://www.thermofisher.com/antibody/product/Donkey-anti-Rabbit-IgG-H-L-Highly-Cross-Adsorbed-Secondary-Antibody-Polyclonal/A-31572 |
| Alexa Fluor 488 | IgG Rat | Donkey | 1:1000 | Invitrogen | A-11006 | RRID:AB_2534074 | https://www.thermofisher.com/antibody/product/Goat-anti-Rat-IgG-H-L-Cross-Adsorbed-Secondary-Antibody-Polyclonal/A-11006 |
| Alexa Fluor 488 | IgG Chicken | Donkey | 1:1000 | Strattech Scientific Ltd | 703-545-155-JIR-0.5mg | RRID:AB_2340375 | https://www.jacksonimmuno.com/catalog/products/703-545-155 |
| DAPI (4',6-Diamidino-2-Phenylindole, Dilactate) | DAPI | NA | 1:1000 | Thermo Fisher Scientific | D1306 | RRID:AB_2307445<br>or<br>RRID:AB_2629482 | https://www.thermofisher.com/order/catalog/product/D1306 |
| Streptavidin, Alexa Fluor™ 488 Conjugate | Neurobiotin | Donkey | 1:1000 | Thermo Fisher Scientific | S32354 |  | https://www.thermofisher.com/order/catalog/product/S32354 |

## Data Availability

The original data used in this study is available at Zenodo (DOI 10.5281/zenodo.7661355), and detailed protocols of each method is available at protocols.io (DOI 10.17504/protocols.io.x54v9dw61g3e/v1).

## Author Contributions

Q.D. designed, performed, and analysed all experimental data. N.I.A. and C.L. undertook iPSC-dopaminergic neuron differentiations. B.N. optimised iPSC-cortical neuron differentiation. J.B. provided expertise and assistance with fabrication of microfluidic chambers. N.B.V. and Q.D. fabricated microfluidic chambers. N.B.V. and R.W.M conceived the project. N.B.V., R.W.M and R.M.G. supervised the study. Q.D., N.B.V. and R.W.M. wrote the paper, with contributions from all authors. All authors reviewed the manuscript and approved its submission.

## Acknowledgements

This research was funded in part by Aligning Science Across Parkinson’s [ASAP-020370] through the Michael J. Fox Foundation for Parkinson’s Research (MJFF), and in part by the Monument Trust Discovery Award from Parkinson’s UK (J-1403). The work was supported by a National Institute for Health Research-Medical Research Council Dementias Platform UK Equipment Award (MR/M024962/1) to R.W.M. Q.D. was supported by a National Science Scholarship from Agency for Science, Technology and Research in Singapore. For the purpose of open access, the author has applied a CC BY public copyright license to all Author Accepted Manuscripts arising from this submission. We thank Natalie Connor-Robson for her advice on MSN differentiation.

---

**Supplementary figure 1:**
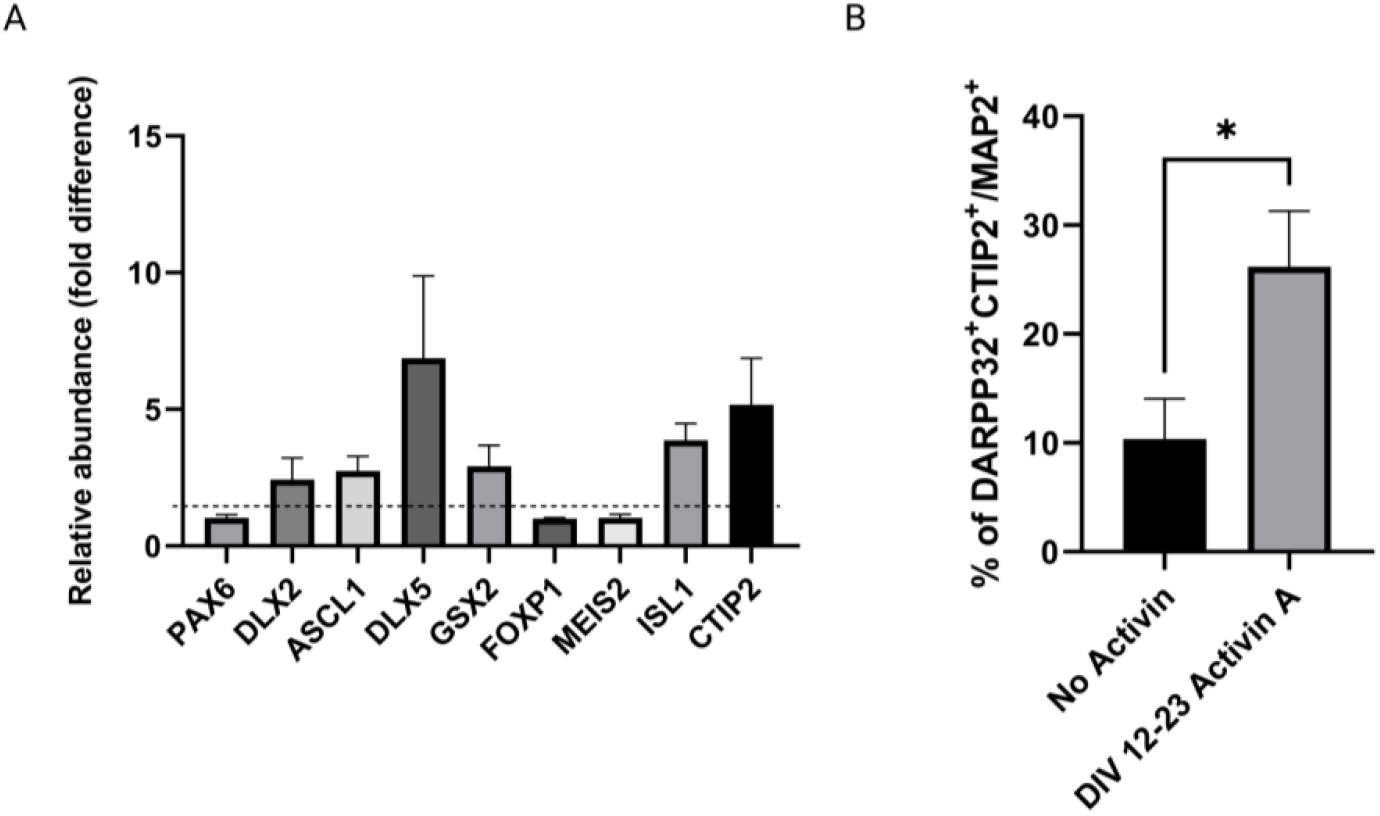
Activin A promoted striatal development. (A) Relative transcript abundance of selected LGE and MSN progenitor markers at DIV14 when cells were treated with activin A at DIV 12 normalised against those that were not; dotted line represents fold difference of 1. (B) Percentages of post-mitotic neurons co-expressing DARPP32 and CTIP2 at DIV 40 in the presence or absence of Activin A from DIV 12-23. N = 2-3 differentiation, n = 3-4 iPS cell lines, and 3 technical replicates.

**Supplementary figure 2:**
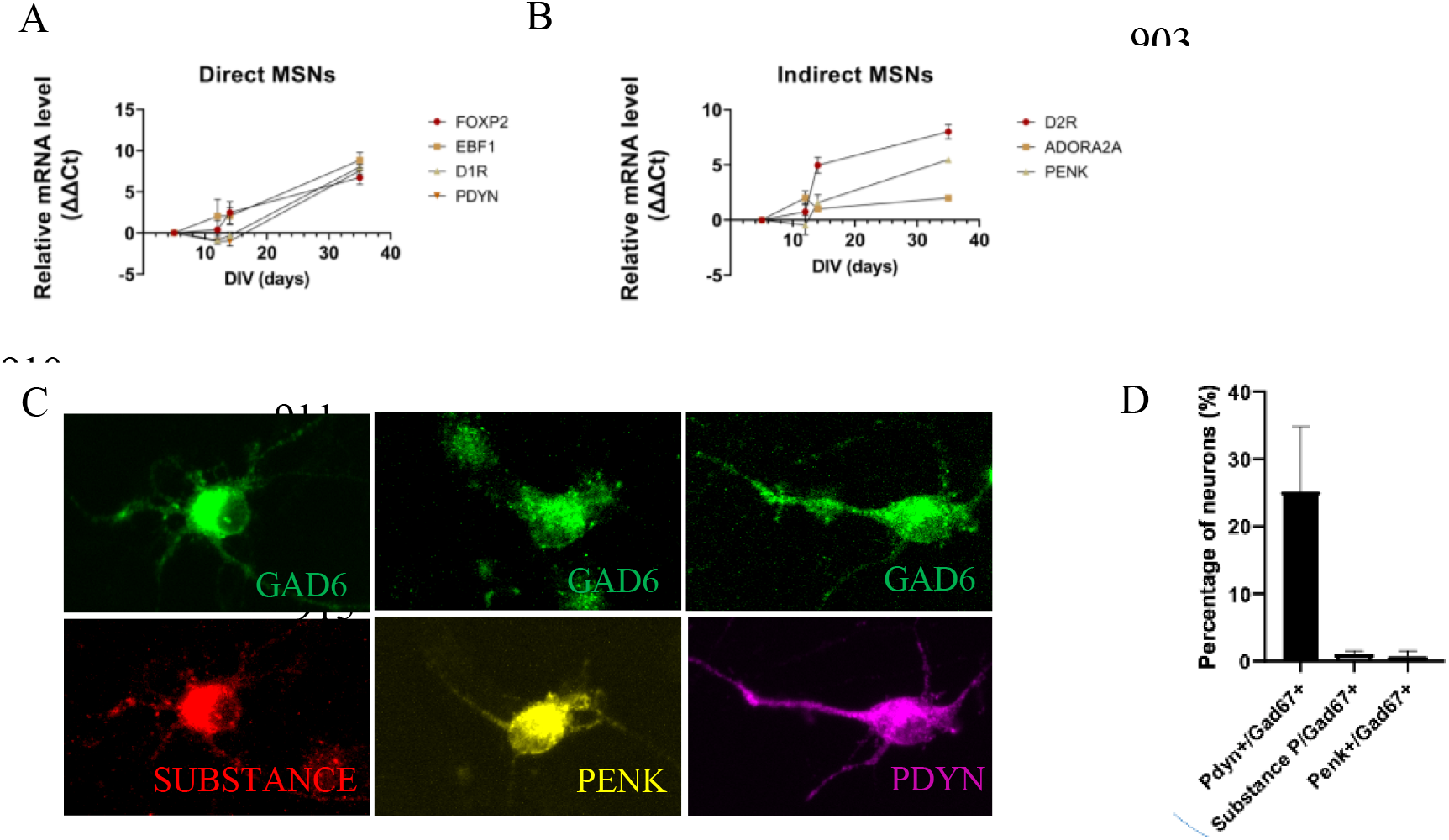
Cultures contained a mixture of both direct and indirect MSNs. (A-B) Transcript abundance of selected marker of (A) direct MSNs and (B) indirect MSNs over time. Expression was averaged from 2 differentiations of 3 independent iPS cell lines. (C) Immunostaining for GAD67 (green), SUBSTANCE P (red), PKEN (yellow), and PDYN (magenta) at DIV 50 (D) Proportion of GAD67+ cell co-expresses different MSN subtype-specific neurotransmitter at DIV 50. All data is presented as mean ± sem, N = 2 differentiation experiments of 3 iPS cell lines.

**Supplementary Figure 3:**
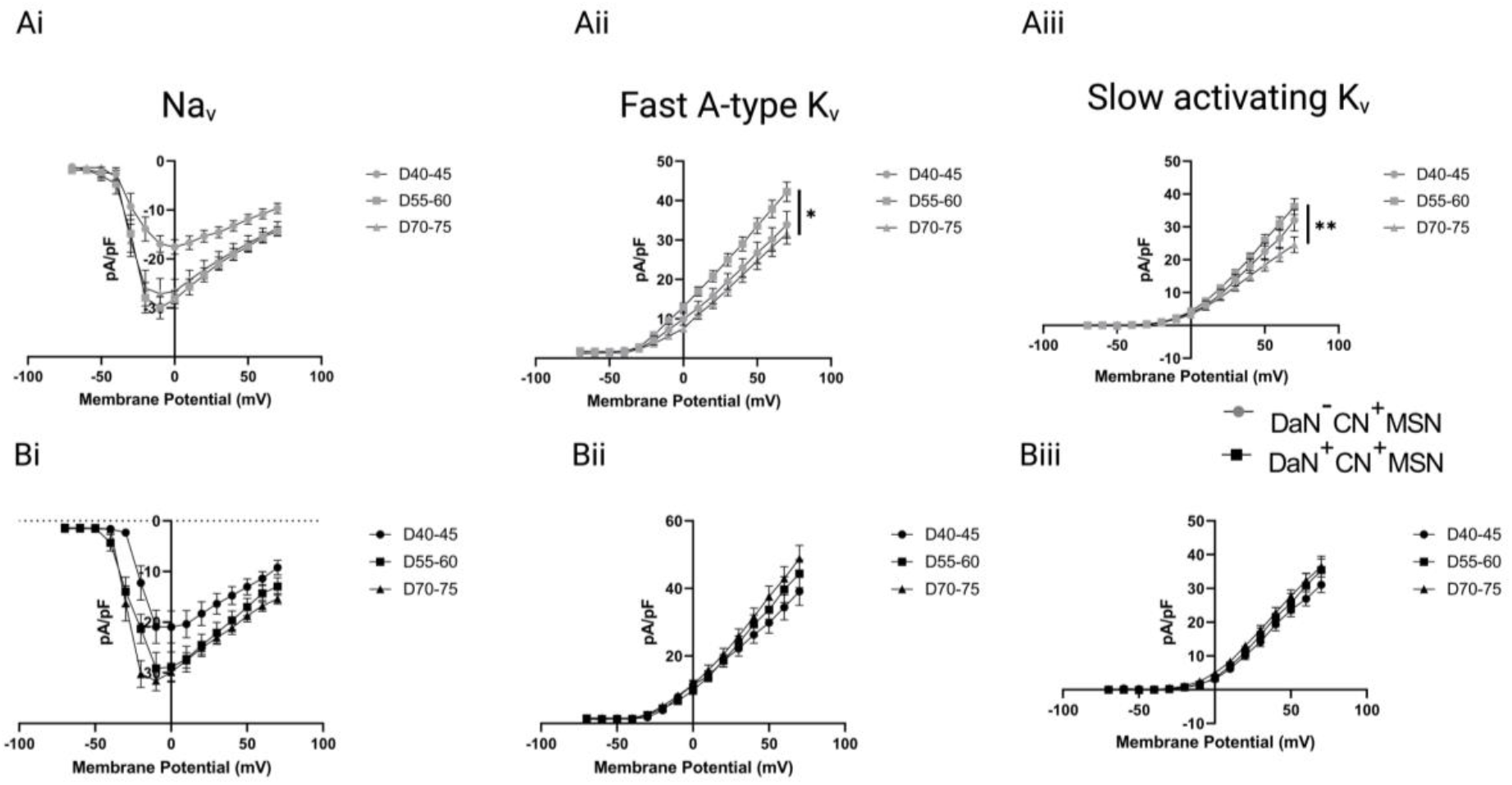
Dopaminergic transmission mediated temporal changes of K_v_ current density. (A-B) Temporal profile of sodium and potassium ionic currents of cortico-striatal neurons in the (A) present and (B) absence of dopaminergic input.

